# Conserved Keratin Gene Clusters in Ancient Fish: an Evolutionary Seed for Terrestrial Adaptation

**DOI:** 10.1101/2020.05.06.063123

**Authors:** Yuki Kimura, Masato Nikaido

## Abstract

Type I and type II keratins are subgroups of intermediate filament proteins that provide toughness to the epidermis and protect it from water loss. In terrestrial vertebrates, the keratin genes form two major clusters, clusters 1 and 2, each of which is dominated by type I and II keratin genes. By contrast, such clusters are not observed in teleost fish. Although the diversification of keratins is believed to have made a substantial contribution to terrestrial adaptation, its evolutionary process has not been clarified. Here, we performed a comprehensive genomic survey of the keratin genes of a broad range of vertebrates. As a result, we found that ancient fish lineages such as elephant shark, reedfish, spotted gar, and coelacanth share both keratin gene clusters. We also discovered an expansion of keratin genes that form a novel subcluster in reedfish. Syntenic and phylogenetic analyses revealed that two pairs of *krt18/krt8* keratin genes were shared among all vertebrates, thus implying that they encode ancestral type I and II keratin protein sets. We further revealed that distinct keratin gene subclusters, which show specific expressions in the epidermis of adult amphibians, stemmed from canonical keratin genes in non-terrestrial ancestors. Molecular evolutionary analyses suggested that the selective constraints were relaxed in the adult epidermal subclusters of amphibians as well as the novel subcluster of reedfish. The results of the present study represent the process of diversification of keratins through a series of gene duplications that could have facilitated the terrestrial adaptation of vertebrates.

**Highlights:**

- Two major keratin clusters are conserved from sharks to terrestrial vertebrates.
- Adult epidermis-specific keratins in amphibians stem from the two major clusters.
- A novel keratin gene subcluster was found in reedfish.
- Ancestral *krt18/krt8* gene sets were found in all vertebrates.
- Functional diversification signatures were found in reedfish and amphibian keratins.

## 1. Introduction

Vertebrate skin structure dramatically changed during the evolutionary transition of vertebrates from aquatic living as fish to terrestrial living as mammals. In the process of terrestrial adaptation, amniotes (e.g., mammals, birds, and reptiles) acquired the stratum corneum, an outer epidermal layer that is filled with keratinous fibers, to guard the skin against drying (Schempp et al., 2009; Yokoyama et al., 2018). By contrast, the skin of bony fish is covered with bony scales to protect them from predators. Amphibians, a basal lineage of terrestrial vertebrates, do not possess scales. According to Szarski (1962), the lack of scales allows amphibians to reduce their body weight as well as to breathe through their skin. Alibardi (2001) described the existence of an amniote-like stratum corneum in the epidermis of adult frogs, but not in tadpoles or aquatic lungfish, implying that the acquisition of the stratum corneum is key to terrestrial adaptation.

Keratin proteins are intermediate filaments and main components of the stratum corneum. Keratin proteins combine with filaggrin to provide durability and prevent water loss from epidermal cells (Sandilands et al., 2009). Keratin proteins are divided into two classes (type I and type II), which are strictly interdependent for assembly into filaments and form coiled-coil heterodimers during the first stage of the process (Jacob et al., 2018). In tetrapods, keratin genes form two clusters (clusters 1 and 2), each of which is mostly dominated by type I and II keratin genes, respectively (Hesse et al., 2004; Zimek and Weber, 2006). However, such gene clusters were not observed in teleost fishes (Zimek and Weber, 2005; Vandebergh and Bossuyt, 2012). Phylogenetic analyses revealed the reciprocal monophyly of type I and type II keratin genes (Schaffeld and Schultess, 2006; Suzuki et al., 2017). The expansion of the keratin genes is believed to have contributed to the terrestrial adaptation (Vandebergh and Bossuyt, 2012). For example, terrestrial vertebrates possess a higher number of keratin genes compared to teleosts despite the whole genome duplication that teleosts underwent. However, this may be the result of massive gene loss that followed this event (Zimek and Weber, 2005). Aquatic mammals lost several keratin genes for outer epidermal layers that would be essential in order to effectively survive in terrestrial environments (Ehrlich et al., 2019). Some keratin genes show specific expressions in particular developmental stages and tissues in adult epidermis (Watanabe et al., 2001, 2002). A recent genome-wide analysis on keratin genes in *Xenopus laevis* and *X. tropicalis* revealed that they possess 4–10 genes for both type I and II keratins, which are specifically expressed in the adult epidermis. These keratin genes form two “adult epidermal keratin subclusters” that are located in each of the two keratin gene clusters (Suzuki et al., 2017).

Although the expansion of the keratin genes in the clusters may be involved in the terrestrial adaptation of vertebrates, the origin and process of keratin gene diversification still remain unclear. Until now, type I and II keratin genes were characterized not only in terrestrial vertebrates but also in early diverged vertebrates such as lamprey and shark (Vandebergh and Bossuyt, 2012); bichir, sturgeon, and gar (Schaffeld et al., 2007); teleosts (Zimek and Weber, 2005); and lungfish (Schaffeld et al., 2005). The origin of the keratin gene clusters was expected to date back to Chondrichthyes (Vandebergh and Bossuyt, 2012). However, the existence of the cluster has not yet been explored in the genomes of Chondrichthyes, Sarcopterygii, and basal Actinopterygii. Thus, characterization and comparison of the clusters in the species of these classes at the whole genome level are of chief importance in order to fill the evolutionary gap between fish and terrestrial vertebrates. In the present study, we revealed the existence of two major keratin clusters, which are conserved among Chondrichthyes, basal Actinopterygii, coelacanths, and terrestrial vertebrates. We also discovered that a set of *krt18/krt8* genes is highly conserved in all vertebrates in a head-to-head array, implying that they could be ancestral keratin pairs. In addition to these two major clusters, we found a novel subcluster of keratin genes that were specifically expanded in reedfish. Phylogenetic and syntenic analyses revealed that the keratin genes belonging to the “adult epidermal keratin subclusters” stem from canonical keratin genes that have already existed in the keratin clusters of non-terrestrial ancestors. Selection analysis suggests relaxation of purifying selection in the reedfish-specific subcluster and the “adult epidermal keratin subclusters” of amphibians. On the basis of our results, we discuss the evolutionary process and the possible contribution of keratin gene expansion that lead to epidermal diversification and terrestrial adaptation in vertebrates.

## 2. Methods

### 2.1 Keratin gene sequences and locations

We collected data for the sequences, locations, and orientation of keratin genes in *Callorhinchus milii* (elephant shark; Assembly Callorhinchus_milii-6.1.3; NCBI Annotation Release 100; Venkatesh et al., 2014), *Erpetoichthys calabaricus* (reedfish; Assembly fErpCal1.1; NCBI Annotation Release 100), *Lepisosteus oculatus* (spotted gar; Assembly LepOcu1; NCBI Annotation Release 101; Braasch et al., 2016), *Latimeria chalumnae* (coelacanth; Assembly LatCha1; NCBI Annotation Release 101; Amemiya et al., 2013), *Rhinatrema bivittatum* (caecilian; Assembly aRhiBiv1.1; NCBI Annotation Release 100), *Xenopus tropicalis* (western clawed frog; Assembly UCB_Xtro_10.0; NCBI Annotation Release 104; Hellsten et al., 2010) and *Petromyzon marinus* (lamprey; Assembly kPetMar1 .pri; NCBI Annotation Release 100) that were deposited in the National Center for Biotechnology Information (NCBI) Database (https://www.ncbi.nlm.nih.gov/). Data for *Branchiostoma belcheri* (amphioxus) were collected from Branchiostoma.belcheri_HapV2(v7h2) through LanceletDB (http://genome.bucm.edu.cn/lancelet/; You et al., 2019). All of the collected sequences were manually inspected to determine whether the gene was pseudogenized or not. The amino acid and nucleotide sequences, and the location of the intact and pseudogenes were summarized in Supplementary Files S1–3. The keratin gene sequences of *Protopterus dolloi* (lungfish) were obtained from RNA-seq data for skin tissue (SRX4966457) and pectoral fins (SRX895362; Takezaki and Nishihara, 2016). Raw read data were assembled *de novo* with Trinity (Grabherr et al., 2011) using the default settings and were extracted keratin gene sequences from assembled data with FATE (Suzuki, 2017) with BLASTN (Altschul et al., 1990) and exonerate (Slater and Birney, 2005) using the coelacanth keratin gene as a query. If the sequences were more than 99% similar, only one sequence was extracted. Gene names were assigned according to NCBI nomenclature and previous studies.

### 2.2 Phylogenetic analysis for all keratin genes

We constructed a phylogenetic tree of keratin genes for a broad range of vertebrates and amphioxi. Amino acid sequences for keratins of amphioxi, lampreys, elephant sharks, reedfish, spotted gars, coelacanths, lungfish, caecilians, and western clawed frogs were used for analysis. These sequences were aligned using MAFFT version 7 (Katoh and Toh, 2008) with the E-INS-i parameter, and over 50% of gap sites were removed. The phylogenetic tree was generated using RAxML version 8.2.12 (Stamatakis, 2014) with the amino acid substitution LG + F model and 300 bootstrap replicates. The model was determined using MEGA X (Kumar et al., 2018).

### 2.3 Quantitative analysis: expression of epidermal keratin in adult amphibians

To identify keratin genes that are expressed in the epidermis of adult amphibians, we analyzed RNA-seq data *of Rhinatrema bivittatum* skin (SRR5591419; Torres-Sánchez et al., 2019) and *Xenopus tropicalis* skin (SRR1405691 and SRR1405692). The read data were checked for quality, and quality control was performed with PRINSEQ++ (Cantu et al., 2019) using the following settings: “-trim_left 5 -trim_tail_right 5 -trim_qual_right 30 -ns_max_n 20 -min_len 30.” After quality control, the read data were mapped to reference genomes (aRhiBiv1.1 and UCB_Xtro_10.0) with STAR version 2.7 (Dobin et al., 2013), and the read counts were normalized with TPMCalculator (Vera Alvarez et al., 2019). The resulting data were summarized in Supplementary Table S2. TPM values (>100) were used as criteria to determine whether keratin genes were expressed in the adult epidermis.

### 2.4 Selection analysis

To evaluate the relaxation of purifying selection on sequences of keratin genes, we calculated the non-synonymous substitution rate to synonymous substitution rate ratio (dN/dS) for several clades of interest. The nucleotide sequences of keratin genes were analyzed by separating them into type I and type II. The translated amino acid sequences were aligned in MAFFT using default parameters (Katoh et al., 2002). This multiple alignment of amino acid sequences was converted into a codon alignment using PAL2NAL (Suyama et al. 2006). We used the CodeML program in PAML 4.9j (Yang, 2007) to analyze branch models (Yang, 1998) for branches where the keratin genes revealed an expansion in copy numbers. We used the CF2 model, where codon frequencies are calculated from the average nucleotide frequencies at the three codon positions, for analysis. To assess the statistical significance of the elevation of the *d*_N_/*d*_S_ ratio for each branch, likelihood ratio tests and inspections of the p-value were used to compare likelihoods between two models by assuming that the *d*_N_/*d*_S_ ratio was not changed in all branches (null hypothesis) and that the *d*_N_/*d*_S_ ratio was only changed in foreground branches (alternative hypothesis).

## 3. Results

### 3.1 Keratin gene clusters conserved among ancient fishes

An exploration of keratin genes from the genomes from one Cephalochordata (amphioxus), one Cyclostomata (sea lamprey), one Chondrichthyes (elephant shark), two basal Actinopterygii (spotted gar and reedfish), one Actinistia (coelacanth), and two Amphibia (caecilian and western clawed frog) gave an overview of the genomic organization of vertebrate keratin gene clusters (Fig. 1; Supplementary Table S1). In terrestrial vertebrates, type I keratin genes in cluster 1 are flanked by *smarce1* and *eif1*, and type II keratin genes (and only one type I keratin gene) in cluster 2 are flanked by *faim2* and *eif4b*. In teleosts, however, syntenic relationships were not observed in either type I or type II keratin genes (Zimek and Weber, 2005; Vandebergh and Bossuyt, 2012). Here, we revealed that the syntenic relationship of type I keratin genes in cluster 1 was highly conserved among elephant shark, spotted gar, reedfish, coelacanth, and terrestrial tetrapod genomes (Fig. 1). The type I keratin genes of reedfish and spotted gar that are flanked by *smarce1* and *eif1* in cluster 1 were monophyletic in the phylogenetic tree (Fig. 2; D). We named these keratin genes “*krt49*.” One of the *krt49* genes – *krt49.1* – exists outside the cluster that is bounded by *smarce1*.

**Fig. 1.**
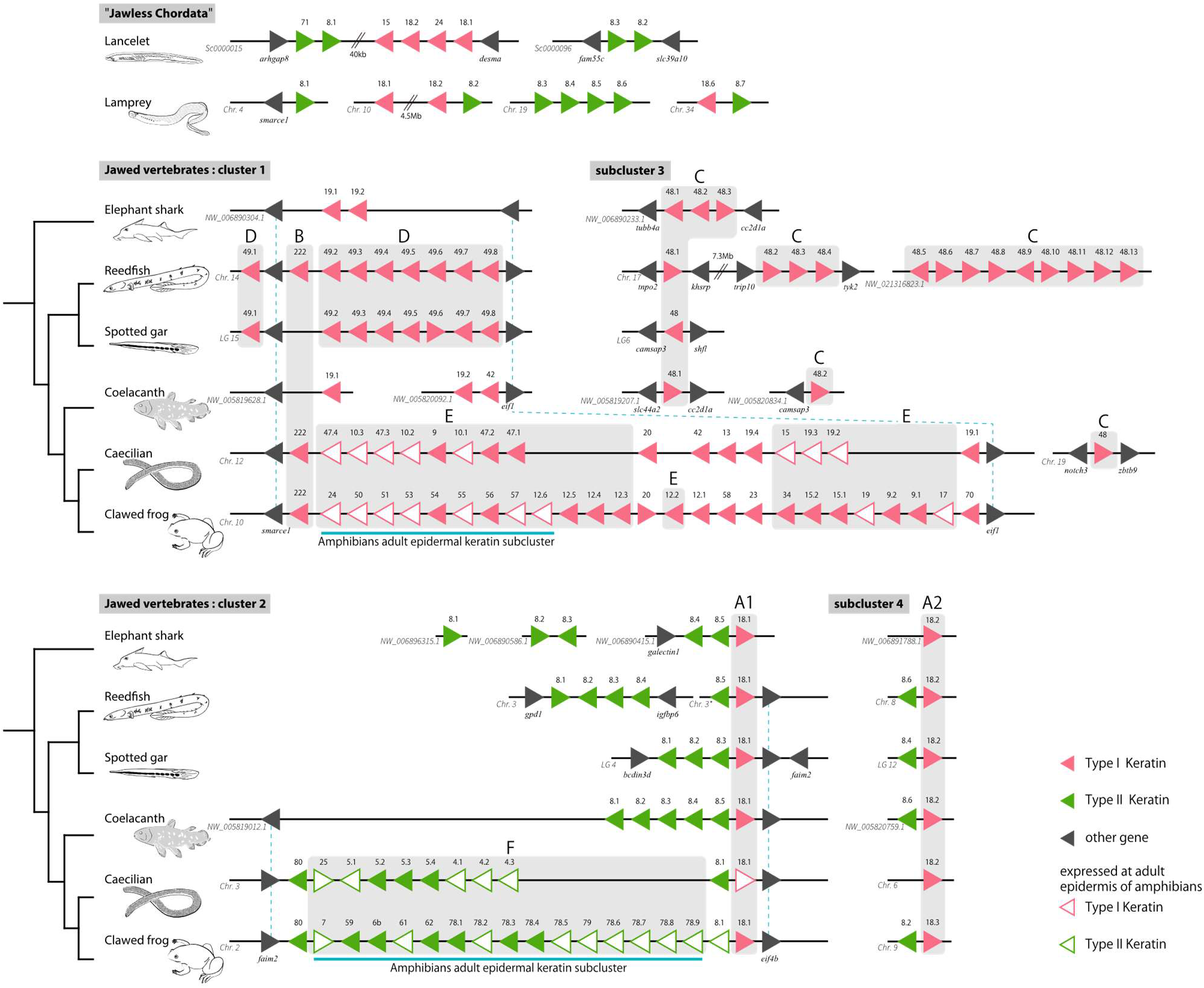
Comparison of keratin gene clusters in vertebrates and amphioxus. The keratin gene clusters were compared among amphioxus (*Branchiostoma belcheri*), lamprey (*Petromyzon marinus*), elephant shark (*Callorhinchus milii*), reedfish (*Erpetoichthys calabaricus*), spotted gar (*Lepisosteus oculatus*), coelacanth (*Latimeria chalumnae*), caecilian (*Rhinatrema bivittatum*), and western clawed frog (*Xenopus tropicalis*). Lines indicate a continuous genomic scaffold, linkage group, or chromosome. Double slashes indicate an abbreviated genomic region. Type I keratin genes, type II keratin genes, and non-keratin genes are indicated by pink, green, and black triangles, respectively. The keratins that are expressed in the epidermis of adult amphibians are indicated with white triangles. All triangles indicate the transcriptional direction of genes for each chromosome or scaffold (LG). Gray boxes with letters correspond to the clades in the phylogenetic tree (Fig. 2). The chromosome, linkage group, and scaffold are shown in the lower left of black lines. Asterisk: unlocalized scaffolds of chromosome 3.

**Fig. 2.**
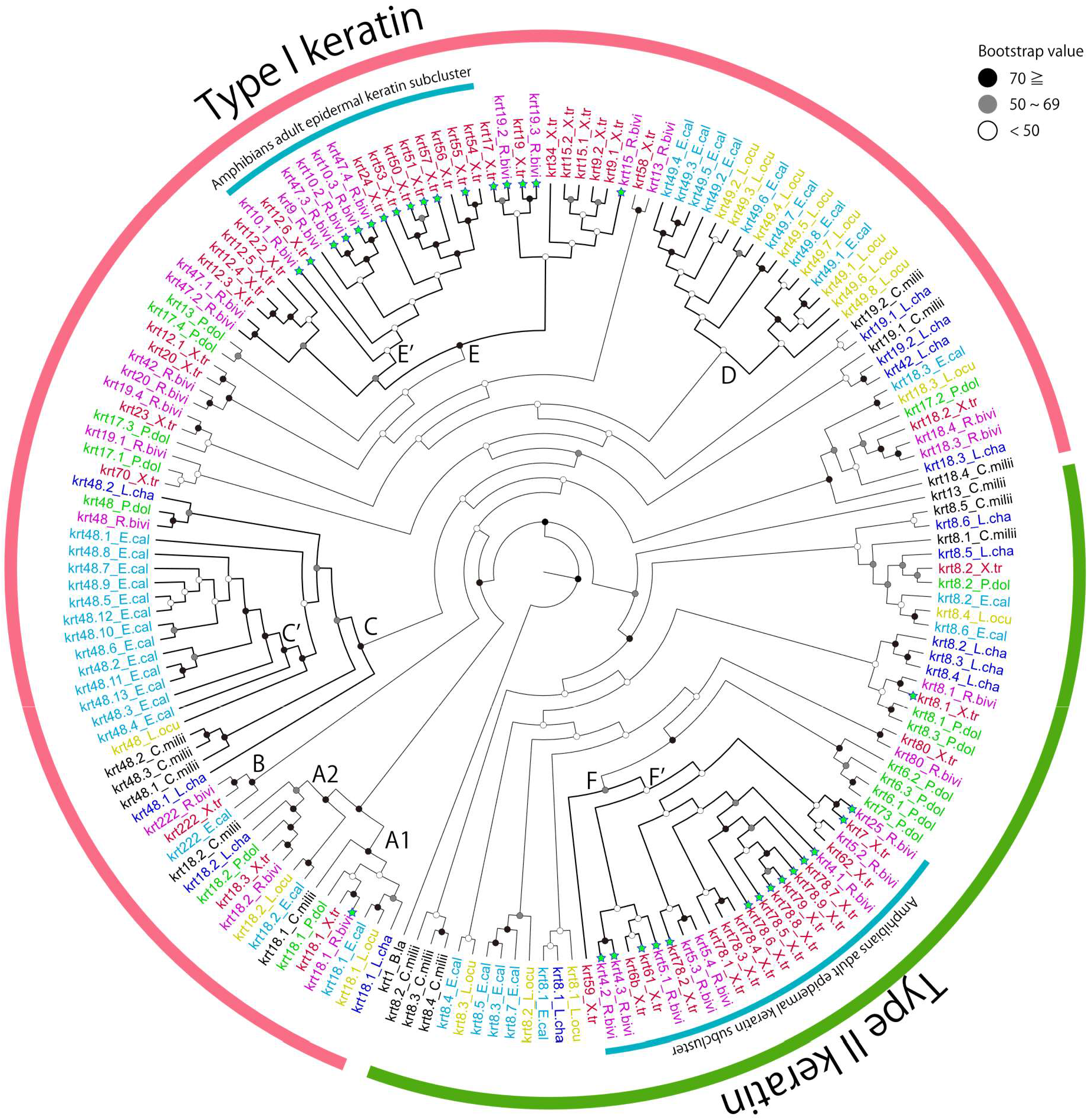
Phylogenetic tree of type I and II keratin genes. The species included in the tree are the following: elephant shark (C. *milii*; C.milli, black), reedfish (E. *calabaricus*; E.cal, light blue), spotted gar (L. *oculatus*; L.ocu, yellow), coelacanth (L. *chalumnae*; L.cha, blue), lungfish (*P. dolloi*; P.dol, green), caecilian (R. *bivittatum*; R.bivi, purple), and western clawed frog (*X. tropicalis*; X.tr, red). The sequence of keratin k1 of *Branchiostoma lanceolatum* was designated as the outgroup (CAB75942.1). Black, gray, and white circles on the nodes indicate bootstrap values 70 and over, 50 to 69, and under 50, respectively. Green stars indicate the expression of keratin genes in the skin of adult *R. bivittatum* and *X. tropicalis*. Letters near the nodes correspond to the gray boxes in Fig. 1.

We also found syntenic relationships in type II keratin genes in cluster 2, although they were somewhat weak compared to those in cluster 1. For example, coelacanths possess the type II keratin gene cluster, which is flanked by *eif4b* and *faim2*, as shown in terrestrial tetrapods. In spotted gars and reedfish, *eif4b* exists at the end of the cluster, but *faim2* is absent in the opposite end of the cluster. Elephant sharks possess cluster 2, which is represented by several type II genes but only one type I keratin gene (*krt18*s). However, it was not flanked by *eif4b* and *faim2* (Fig. 1).

In addition to the major clusters 1 and 2 localized in two genomic regions (Vandebergh and Bossuyt, 2012), we found a novel keratin gene subcluster 3, where 13 genes were tandemly duplicated, in reedfish (Fig. 1 C). The keratin genes arranged in this subcluster are homologs of *krt48s*. Initially, *krt48s* were identified by Schaffeld et al. (2007) as “*krt14*”. We re-named these genes as “*krt48*” to avoid confusion in referring *krt14* in mammals. *krt48s* are shared by elephant sharks, gars, sturgeon, and coelacanths, but were not extensively expanded in these species. None of the *krt48* genes shared any synteny with cluster 1 or 2, implying that this subcluster is distinct from the two major clusters. Interestingly, one copy of *krt48* was found in the basal lineage of the amphibian *Rhinatrema bivittatum* but not in the other terrestrial vertebrates (Figs. 1 and 2). We also found subcluster 4, in which only one set of *krt18s* and *krt8s* existed (Fig. 1). This subcluster is also distinct from the two major clusters 1 and 2 as no synteny relationships were observed. Subcluster 4 exists in a broad range of vertebrates, but it was not found in some terrestrial vertebrates such as caecilians, birds, and mammals.

### 3.2 Highly conserved keratin gene set *krt18/krt8*

We revealed that several keratin genes are highly conserved among a broad range of vertebrates as represented by *krt18*s (clades A1 and A2) and *krt8*s (Fig. 1), which were shown to be shared in terrestrial vertebrates (Balmer et al., 2017; Vandebergh and Bossuyt, 2012; Ehrlich et al., 2019). It is important to note that the head-to-head array of *krt18s* and *krt8s* in cluster 2 and that in subcluster 4, in which the 5’ ends of two adjacent genes face each other, were highly conserved from sharks to terrestrial vertebrates (Fig. 1). The characteristic head-to-head arrays of *krt18s* and *krt8s* are observed exclusively in these two loci. Although the synteny relationships are not obviously identified, we found two pairs of head-to-head *krt18/krt8* in the lamprey genome, the most basal lineage of vertebrates (Fig. 1). We also found the keratin gene sequences, which were assigned as “*krt18s*,” in the amphioxus genome. However, these genes are not arranged in a head-to-head manner with type II keratins (Fig. 1). Phylogenetic analysis suggests that *krt18.1* (clade A1) and *krt18.2* (clade A2) from shark to terrestrial vertebrates are monophyletic and these two clades are most ancestral in type I keratin genes (Fig. 2). The additional phylogenetic analyses that focus on the *krt18*s and *krt8*s of lampreys and amphioxi revealed that the *krt18.1* and *krt18.2* of lampreys are sister groups of clades A1 and A2 (Fig. S1). By contrast, *krt18s* of amphioxi diverged quite earlier than those of vertebrates (Fig. S1). *krt222* (clade B) is also found in a broad range of vertebrates from reedfish to western clawed frogs as well as mammals (Ehrlich et al., 2019). However, they became pseudogenes in elephant sharks, spotted gars, and coelacanths (Fig. 1; B).

### 3.3 Copy number of keratin genes among vertebrates

We counted and compared the total number of intact keratin genes among vertebrates (Fig. 3). By taking the phylogenetic tree into account, the type I keratin genes had increased after the emergence of tetrapods. The number of type I keratin genes was constant in most fish of Actinopterygii lineages, except for reedfish where 25 genes were observed. The increase of type I keratin genes in reedfish is explained by the expansion of *krt48* in subcluster 3 (Fig. 1). The number of type II keratin genes was also constant in fish lineages but suddenly increased in the common ancestor of terrestrial vertebrates. In spite of the fact that the ancestor of teleosts experienced whole genome duplication, the copy number of keratin genes is not higher than that of other fish species. The marked increase of keratin genes was observed in tetrapods and reedfish.

**Fig. 3.**
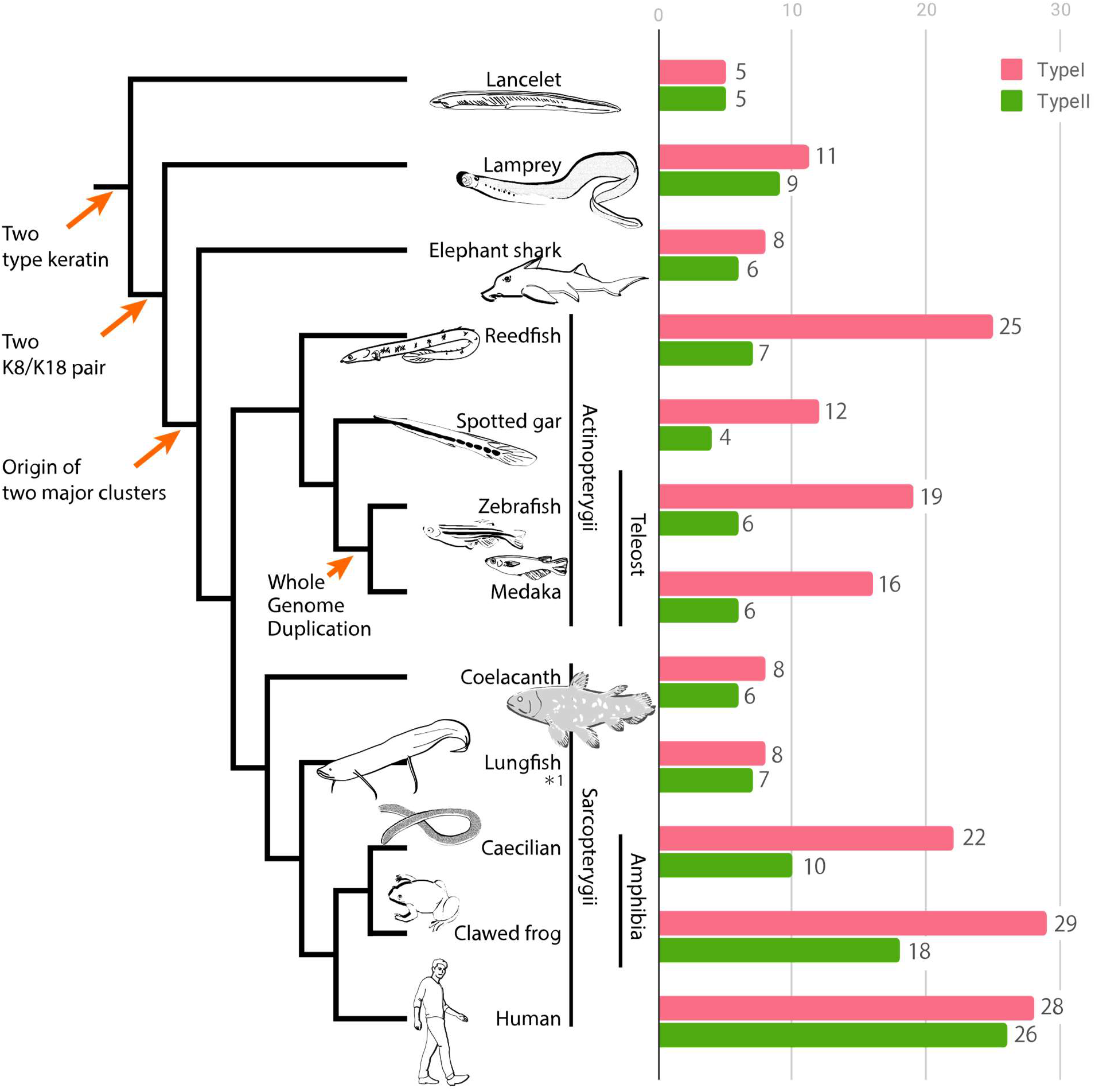
The number of keratin genes among vertebrates and amphioxus. The number of intact keratin genes was compared among Cephalochordata (amphioxus), Cyclostomata (lamprey), Chondrichthyes (Elephant shark), Actinopterygii, and Sarcopterygii (including Amphibia). Type I keratin: pink bar; type II keratin: green bar. Arrows in the phylogenetic tree indicate the timing of the evolutionary events. *1: The number of keratin genes of lungfish estimated using data obtained from RNA-seq analysis.

In addition, we counted the number of cysteine residues in keratins (Fig. S2). The number of cysteines were significantly higher in a mammal and amphibians than in fish, which possess only a few cysteines. Although it is not statistically significant, the number of cysteines were high in lamprey and lungfish.

### 3.4 Origin of “adult epidermal keratin subcluster” in amphibians

In clawed frogs, some keratin genes in both types I and II are specifically expressed in the epidermis of the adult stage but not in the tadpole epidermis (Suzuki et al., 2017; Fig. 2, white triangle). These keratin genes possibly contribute to the adaptation to the terrestrial environment. To explore the evolutionary origin of the adult epidermal keratin subcluster, we performed phylogenetic analyses of keratin genes by adding the keratin genes of ancient fishes and another amphibian species, *Rhinatrema bivittatum*, which is the most basal lineage of extant amphibia (caecilians or Gymnophiona). We also analyzed public RNA-seq data for adult caecilian and western clawed frog to identify which keratin genes were expressed in their skin (Supplementary Table S2). The results of our phylogenetic analysis show that keratin genes, which are expressed in epidermis of caecilians, are included in the clades of the “adult epidermal keratin subcluster” of western clawed frogs (Fig. 2; E, F). Additionally, the locations of these adult epidermal keratin genes in the caecilian genome are close to those in the western clawed frog (Fig. 1; E, F). For lungfish, we also identified the keratin genes that are expressed in skin and pectoral fin using public RNA-seq data. Subsequent phylogenetic analyses suggested that keratin genes expressed in the skin and pectoral fins of lungfish were not included in the amphibian adult epidermal keratin subclusters. It is noteworthy that the phylogenetic and syntenic analyses revealed that the adult epidermal keratins, which belong to clades E and F, shared common ancestry with *krt49s* (clade D, type I) and *krt8s* (type II) that had already existed in the common ancestor of bony fish.

### 3.5 Relaxation of purifying selection in keratin genes

The *d*_N_/*d*_S_ ratio provides useful insight into the strength and mode of natural selection acting on protein-coding genes. In particular, the elevation of the *d*_N_/*d*_S_ ratio can be used to evaluate the operation of natural selection toward functional diversification. To examine the extent of diversification in several keratin gene clusters, we calculated *d*_N_/*d*_S_ values by defining the branches of the clusters of interest as foreground and the other branches as background. We performed the analyses on clades C–F and C’, E’, and F’ in the phylogenetic tree shown in Fig. 2. The copy number of reedfish *krt48* was increased by gene duplication compared to other species (C and C’ in Fig. 2). Clades E and F contain “adult epidermal keratin” subcluster genes of amphibians (Fig. 2). E’ and F’ were monophyletic groups of amphibian adult epidermal keratin subcluster genes. The summary of this analysis is shown in Table 1. The d_N_/d_S_ values among all branches were within the range of 0.1 to 0.3, suggesting that most keratin genes may be under purifying selection. Importantly, however, the d_N_/d_S_ values of C’ (reedfish-specific type I *krt48* branch), E’, and F’ (type I and type II amphibian “adult epidermal keratin” subcluster branches) were higher (C’: 0.29, E’: 0.25, and F’: 0.15) than background (0.18, 0.18, and 0.13, respectively) with statistical significance. The results suggest that the constraints of purifying selection were relaxed in these branches, implying the operation of natural selection toward functional diversification in these clusters.

**Table 1.** Summary of the results of selection analyses and likelihood ratio tests.

| clade C | np<sup>1</sup> | lnL | p-value<sup>2</sup> | $d_N/d_S$ |
| --- | --- | --- | --- | --- |
| Null hypothesis | 224 | −65270.5166 | 0.0679 | 0.1797 |
| Alternative hypothesis | 225 | −65268.8495 |  | 0.2066 |
| clade C' | np | lnL | p-value | $d_N/d_S$ |
| Null hypothesis | 224 | −65270.5166 | 0.0002 | 0.1791 |
| Alternative hypothesis | 225 | −65263.7851 |  | 0.2919 |
| clade D | np | lnL | p-value | $d_N/d_S$ |
| Null hypothesis | 224 | −65270.5166 | 0.1937 | 0.1796 |
| Alternative hypothesis | 225 | −65269.6720 |  | 0.1951 |
| clade E | np | lnL | p-value | $d_N/d_S$ |
| Null hypothesis | 224 | −65270.5166 | 0.1145 | 0.1772 |
| Alternative hypothesis | 225 | −65269.2711 |  | 0.1916 |
| clade E' | np | lnL | p-value | $d_N/d_S$ |
| Null hypothesis | 224 | −65270.5166 | 4.E−10 | 0.1702 |
| Alternative hypothesis | 225 | −65250.8822 |  | 0.2525 |
| clade F | np | lnL | p-value | $d_N/d_S$ |
| Null hypothesis | 116 | −26448.0867 | 0.0501 | 0.1265 |
| Alternative hypothesis | 117 | −26446.1671 |  | 0.1474 |
| clade F' | np | lnL | p-value | $d_N/d_S$ |
| Null hypothesis | 116 | −26448.0867 | 0.0137 | 0.1250 |
| Alternative hypothesis | 117 | −26445.0481 |  | 0.1518 |

<sup>1</sup>np: number of free parameters. <sup>2</sup>p-value: statistical significance for the likelihood ratio test between null hypothesis and alternative hypothesis.

## 4. Discussion

### 4.1 Conservation of two major keratin gene clusters in early vertebrates

Two keratin gene clusters exist in terrestrial vertebrates, but not in teleost fishes (Zimek and Weber, 2005; Vandebergh and Bossuyt, 2012). Since the existence of keratin gene clusters has not been explored in an ancestral group of vertebrates, the timing for the first appearance of these clusters was unclear. Indeed, type I and II keratin genes were identified in ancient lineages of vertebrates such as lamprey, shark, bichir, sturgeon, and gar (Schaffeld et al., 2007; Vandebergh and Bossuyt, 2012). However, the synteny relationships of these genes could not be uncovered because the genome sequences were not available. In the present study, by exploring the whole genome sequences of the aforementioned species, we revealed the existence of both keratin gene clusters 1 and 2, each of which is flanked by two marker genes (Fig. 1). Although the marker genes were not found in cluster 2 of elephant shark and lamprey, the tandem array of the keratin genes indicates the existence of the cluster in these species. Thus, our present study suggests that the origin of the two keratin gene clusters dates back to the common ancestor of Vertebrata.

Although some ancient fish with keratin cluster (such as reedfish and spotted gar) have ganoid scale, the epidermal skin structures are basically similar with the other teleost fishes (Sire, 1995). Thus, it is unlikely that the presence of keratin cluster itself directly contributed to the acquisition of skin structures adapted to terrestrial environments. The evolutionary significance for the existence of keratin cluster in ancient fish could be explained by “exaptation”. Keratin genes of terrestrial tetrapods are packaged together at particular genomic locations and expression stages (Ehrlich et al., 2019; Suzuki et al., 2017), which may be important to accomplish the terrestrial adaptation. Indeed, recent study revealed that a single enhancer regulates the expression of many beta-keratin genes in the cluster (Liang et al., 2020). From that perspective, the presence of keratin clusters in fish prior to terrestrial adaptation may be an “exaptation”. In other words, the clustered keratin genes arrangement in ancient fish were subsequently “co-opted” in terrestrial tetrapods.

### 4.2 Ancient and conservative keratin gene pair *krt18/krt8*

One of the interesting findings in keratin gene cluster 2 is that *krt18.1* and *krt8s* were located in a head-to-head array that is highly conserved from lamprey to tetrapods (Fig. 1; A1). *krt18.1* is exceptional in that it is arranged with type II keratin genes in cluster 2 even though they belong to type I in the phylogenetic tree (Fig. 2). We newly found that *krt18.2* and *krt8*s were located in a head-to-head array in subcluster 4 that is also conserved among a broad range of vertebrates (Fig. 1; A2). It is worth noting that Schaffeld et al. (2005) presumed that *krt18/krt8* found in terrestrial vertebrates has true orthologs in all Gnathostomata (i.e., jawed vertebrates) and possibly in Cyclostomata (i.e., lamprey and hagfish). Our data support this hypothesis by showing the characteristic synteny relationships of these genes. Phylogenetic analysis suggests that the *krt18*s belonging to A1 and A2, which have head-to-head orientation with *krt8*s, are monophyletic at the basal position in type I keratin genes (Fig. 2). We expect that the two pairs of *krt18*s/*krt8*s found in lamprey are the ancestral genes leading to cluster 2 and subcluster 4 in all extant vertebrates. It is implicative that the set of *krt18/krt8* is shown to be first expressed during mammalian embryogenesis (Duprey et al., 1985). These lines of data and observations imply that highly diversified keratin genes in vertebrates originated from the set of *krt18/krt8*, which possess essential and fundamental developmental functions.

### 4.3 Additional *krt48* subcluster in reedfish

In addition to the two major clusters conserved among Gnathostomata (this study; Vandebergh and Bossuyt, 2012), we newly discovered subcluster C in reedfish (Fig. 1). Given that the *krt48*s in subcluster C are monophyletic in the phylogenetic tree (Fig. 2), all of these genes originated from a single ancestral gene through duplications. Initially, *krt48s* were identified by Schaffeld et al. (2007), in which 1 or 2 copies of *krt48s* were only found in basal Actinopterygii (e.g., bichir, sturgeon, and gar). In this study, we revealed that *krt48*s are shared among various vertebrates from shark to amphibians (e.g., caecilian), but not in reptiles, chickens, or mammals. The distribution of *krt48s* in vertebrate genomes suggests that they were acquired in the common ancestor of Gnathostomata and were lost after the divergence of amphibians. Our phylogenetic analysis suggested that *krt48s* diverged next to *krt18s* (Fig. 2). This branching order is not consistent with a previous study (Schaffeld et al., 2007), in which *krt48*s are the basal in type I keratin genes. We suppose that the increase in the number of sequences of *krt48*s and *krt18*s outgroups for the analysis is due to the availability of the whole genome sequences, which may facilitate a more accurate tree estimation.

### 4.4 Genome duplication and copy number variation

It is well established that teleost fish have undergone teleost-specific genome duplication (3R; Meyer and Van de Peer, 2005). Zimek and Weber (2005) presumed that 3R caused the dispersal of the keratin gene cluster, which led to the breakdown of the syntenic relationships of the clusters between teleosts and terrestrial vertebrates. In the present study, we revealed that both of the keratin gene clusters were conserved in basal Actinopterygii, which have not undergone 3R (Fig. 1), supporting the hypothesis by Zimek and Weber (2005) that the 3R resulted in the breakdown of the clusters in teleosts.

We found that the copy number of keratin genes was constant in both types of keratin genes among fish from shark to coelacanth, except for reedfish. In particular, the copy number was also constant in teleosts despite the 3R that they experienced that led to the disruption of gene clusters (Fig. 3). The conservative copy number among fish implies that these keratins were under strict purifying selection for survival in underwater environments. By contrast, the number of keratin genes was expanded in terrestrial vertebrates and reedfish, which raises the possibility of adaptive evolution in these groups (see latter discussion).

It is noteworthy that the number of type I and II keratins are almost symmetric in mammals, shark and lamprey (Vandebergh and Bossuyt, 2012; Ehrlich et al., 2019; this study), whereas asymmetric in Actinopterygii and amphibians (Fig. 3). One possible explanation for the symmetry in mammals may be “one to one co-expression” of type I and II keratin pairs. However, only a limited number of examples for one to one co-expression were shown in mammals (ex. Chu and Weiss, 2002) as well as in fish. Therefore, at present, it is difficult to provide clear answer for the symmetry/asymmetry of the number of type I and II keratins. We can only to discuss that difference in the number of type I and II keratins in Actinopterygii is due to the acceleration of species-specific duplication of type I genes in this group.

### 4.5 Origin of keratin genes specific to terrestrial vertebrates

Terrestrial vertebrates possess a higher number of keratin genes compared to fish, which may have facilitated their invasion of terrestrial environments. The keratin genes that belong to the “adult epidermal keratin subcluster” in clawed frogs (Suzuki et al., 2017) were also shown to be expressed in the skin of adult caecilians (Fig. 1, Fig. 2; E, F). In the present study, no keratin genes that belong to the “adult epidermal keratin subcluster” was found in the RNA-seq data of the skin and fins of lungfish, implying that they do not possess keratin genes belonging to this subcluster. However, it is possible that the “adult epidermal keratin genes” were not expressed in lungfish skin in underwater environments. Indeed, Heimroth et al. (2018) reported that the epidermis of lungfish showed notable changes and became more compact with flattened keratinocytes in response to experimental terrestrial conditions. Therefore, it would be worthwhile to analyze the RNA-seq data collected from the skin of this terrestrial condition to examine the above possibility. Furthermore, analysis of the whole genome, which is not available at present, may lead to the identification of such keratin genes in lungfish.

Phylogenetic and syntenic analyses supported that the “adult epidermal keratin subcluster” of terrestrial vertebrates stemmed from the keratin genes in clusters 1 and 2 that had already existed in the common ancestor of Actinopterygii and Sarcopterygii (Figs. 1, 2). The terrestrial adaptation could be partly facilitated by the diversification of keratin genes through the co-option of existing keratin genes.

### 4.6 Cysteine residues of keratin genes

Recently, crystal structures of human *krt10/krt1* and *krt14/krt5* had been determined (Bunick and Milstone, 2017). In *krt10/krt1* pair, Cys401^K10^, which is conserved among mammals, was shown to be essential to form heterodimer via disulfide link. The mutation at this residue results in a significantly increased trans-epidermal water loss in mice skin (Guo et al., 2020). The cysteine residue at the corresponding site is also conserved in *krt14* among mammals (Lee et al., 2012). To examine the presence/absence of this conserved cysteine residue among broad range of vertebrates, we aligned type I keratin amino acid sequences including human *krt10* and *krt14*. Alignment of keratin sequences revealed that this cysteine residue only exists in keratins expressed in the adult epidermis of amphibians in addition to mammals (Fig. S3). We found that fish do not possess keratins which possess the cysteine residue at the corresponding site of Cys401^K10^. Therefore, our comparative sequence analyses suggest that this cysteine residue may play an important role in terrestrial adaptation.

Our result showed that the number of cysteines in keratins were significantly higher in terrestrial tetrapods (mammals and amphibians) than in fish, which contain only a few cysteines (Fig. S2). Above data imply that the increase of cysteines in keratins may allow ancestral amphibians to acquire hardness in their scaleless epidermis during evolution (Strnad et al., 2011). Considering together with the conserved cysteine residue in the type I keratins (e.g. Cys401^K10^), we suggest that the number of cysteines in keratins have been increased as a result of terrestrial adaptation. It is noteworthy that the number of cysteines were relatively high in lamprey keratins, which may be utilized for keratinized “teeth” (Alibardi and Segalla, 2011) or for thick epidermis with no scales (Elliott, 2011).

### 4.7 Relaxation of purifying selection and diversification of keratin genes

Generally, functional genes are under purifying selection to keep the original function of the protein. In the case of gene duplication, one of the duplicated copies is expected to become free from purifying selection and can acquire a new function, whereas the other copy retains the original function (Ohno, 1970). The relaxation of purifying selection and functional diversification are often discussed in the multigene family such as the olfactory or pheromone receptor gene family (e.g., Niimura et al., 2014; Grus and Zhang, 2004). The relaxation of purifying selection can be elucidated from the elevation of the *d*_N_/*d*_S_ value, which is caused by the acceleration of non-synonymous substitutions. In the present study, we calculated the *d*_N_/*d*_S_ of keratin genes for several clades of interest (Fig. 2; C’, E’, F’ and Table 1) to examine the relaxation of purifying selection. We found that the *d*_N_/*d*_S_ was elevated in subcluster 3 in reedfish and the “adult epidermal keratin subcluster” in amphibians, implying that the keratin genes were diversified in function. In Vandebergh and Bossuyt (2012), duplication and the expansion of the keratin genes contributed to terrestrial adaptation. Indeed, caecilians and western clawed frogs adapted to terrestrial environments, and reedfish are viable in terrestrial environments (Pace and Gibb, 2011; Sacca and Burggren, 1982). Our analyses suggest that the expansion of the keratin genes and relaxation of purifying selection may have facilitated the epidermal adaptation to the terrestrial environment in these species. In particular, diversification of amphibian adult epidermal keratin subcluster (clade E’ and F’) might be important to terrestrial adaptation. The fish skin consists of three main layers: epidermis, scales and dermis. In contrast, amphibians lost the scales and acquired a stratum corneum, which contains various keratins and is important to prevent skin from water loss (Yokoyama et al., 2018). Thus, we expect that there may be an evolutionary link between the diversification of clade E’ and F’ keratin genes and the acquisition of stratum corneum.

## 5. Conclusion

In the present study, we revealed that two major keratin gene clusters are shared among Cyclostomata, Chondrichthyes, basal Actinopterygii, and Sarcopterygii (including terrestrial vertebrates), but not in teleost fish. The result suggests that two major clusters originated in the common ancestor of vertebrates and were conserved among most vertebrates, except for teleost fish, in which the clusters were scattered in multiple chromosomes because of whole genome duplication (3R). The keratin genes belonging to the adult epidermal subclusters in amphibians originated in the two major clusters that had already existed before the timing of the terrestrial adaptation. The specific cysteine residues conserved among mammalian keratins were also found to be conserved in amphibian keratins, which were specifically expressed in adult epidermis, implying that they may contribute to terrestrial adaptation. We found that the pair of *krt18/krt8* genes in a head-to-head array was conserved in all vertebrates, implying the ancestral and essential function of the keratin pairs in epidermal development. The signatures for the relaxation of purifying selection in keratin gene subclusters in reedfish and amphibians represent the possibility of functional diversification in keratins. Thus, the process of terrestrial adaptation in the vertebrate epidermis may be explained by the series of gene duplications from the ancestral keratin genes inherited from ancient lineages.

## Declaration of Competing Interests

The authors have no conflicts of interest to declare.

## Acknowledgments

We thank Yujiro Kawabe of the Tokyo Institute of Technology for the animal illustration included in Fig. 3. We also thank Zicong Zhang of the Tokyo Institute of Technology for technical assistance during evolutionary analyses.

## Funding Sources

This work was funded by JSPS KAKENHI (17K19422) and an Asahi Glass Foundation grant awarded to M.N.

## Supplementary Data

**Supplementary Fig. S1.**
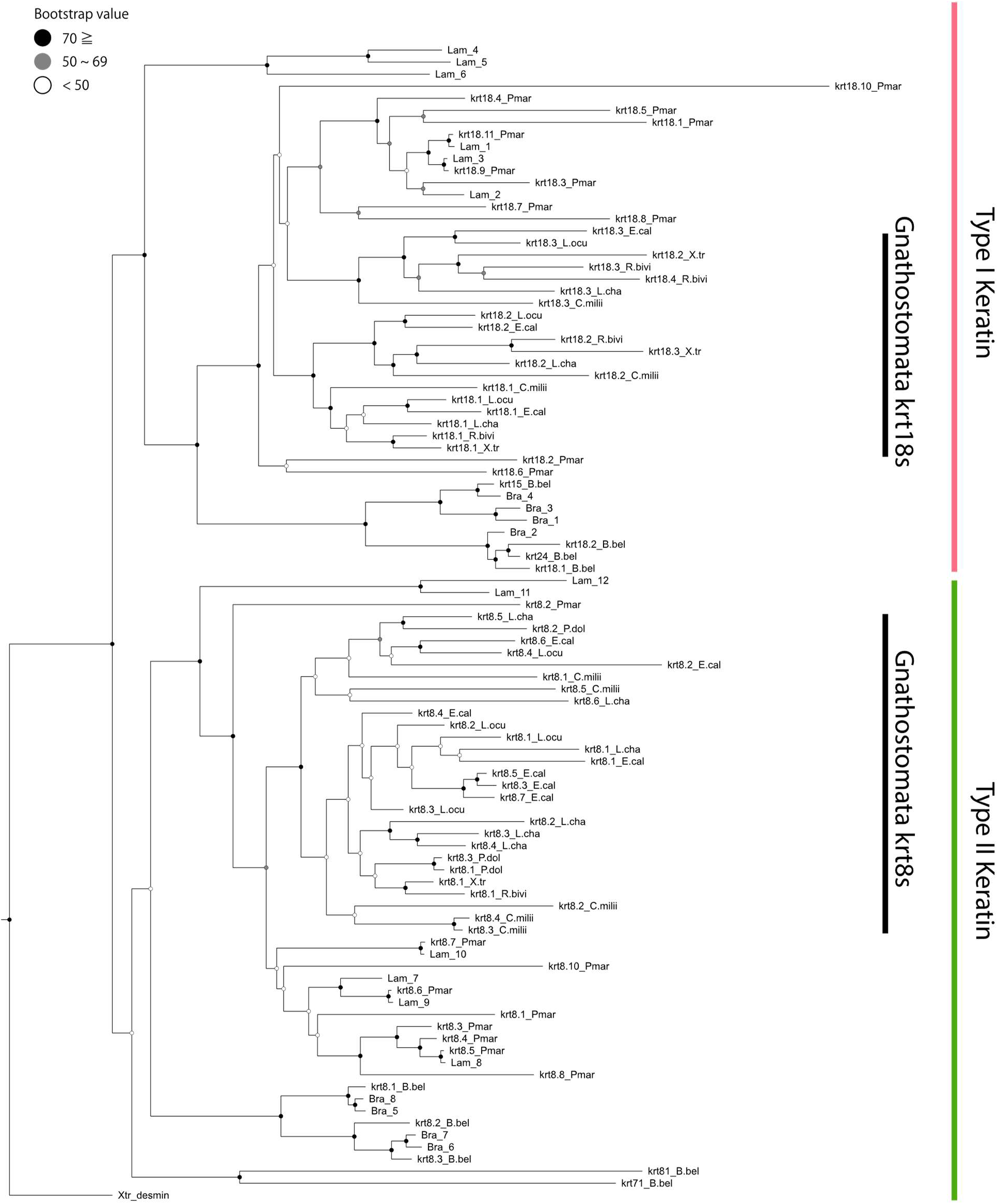
Phylogenetic tree of type I and II keratin genes with focus on the relationships of putative *krt18s* and *krt8s* in amphioxus and lamprey. The species included in the analyses are the following: amphioxus (*Branchiostoma belcheri*; B.bel), lamprey (*Petromyzon marinus*; P.mar), elephant shark (*C. milii*; C.milli), reedfish (*E. calabaricus*; E.cal), spotted gar (*L. oculatus*; L.ocu), coelacanth (L. *chalumnae*; L.cha), lungfish (*P. dolloi*; P.dol), caecilian (*R. bivittatum*; R.bivi), and western clawed frog (*X. tropicalis*; X.tr). We also included the keratin gene sequences of other species of amphioxus (B. *floridae* and *B. lanceolatum*: Bra) and lamprey (*Lampetra fluviatilis*: Lam) that were reported in a previous study (Vandebergh and Bossuyt, 2012). *Desmin* sequence of western clawed frog was used as an outgroup. Black, gray, and white circles on the nodes indicate bootstrap values 70 and over, 50 to 69, and under 50, respectively.

**Supplementary Fig. S2.**
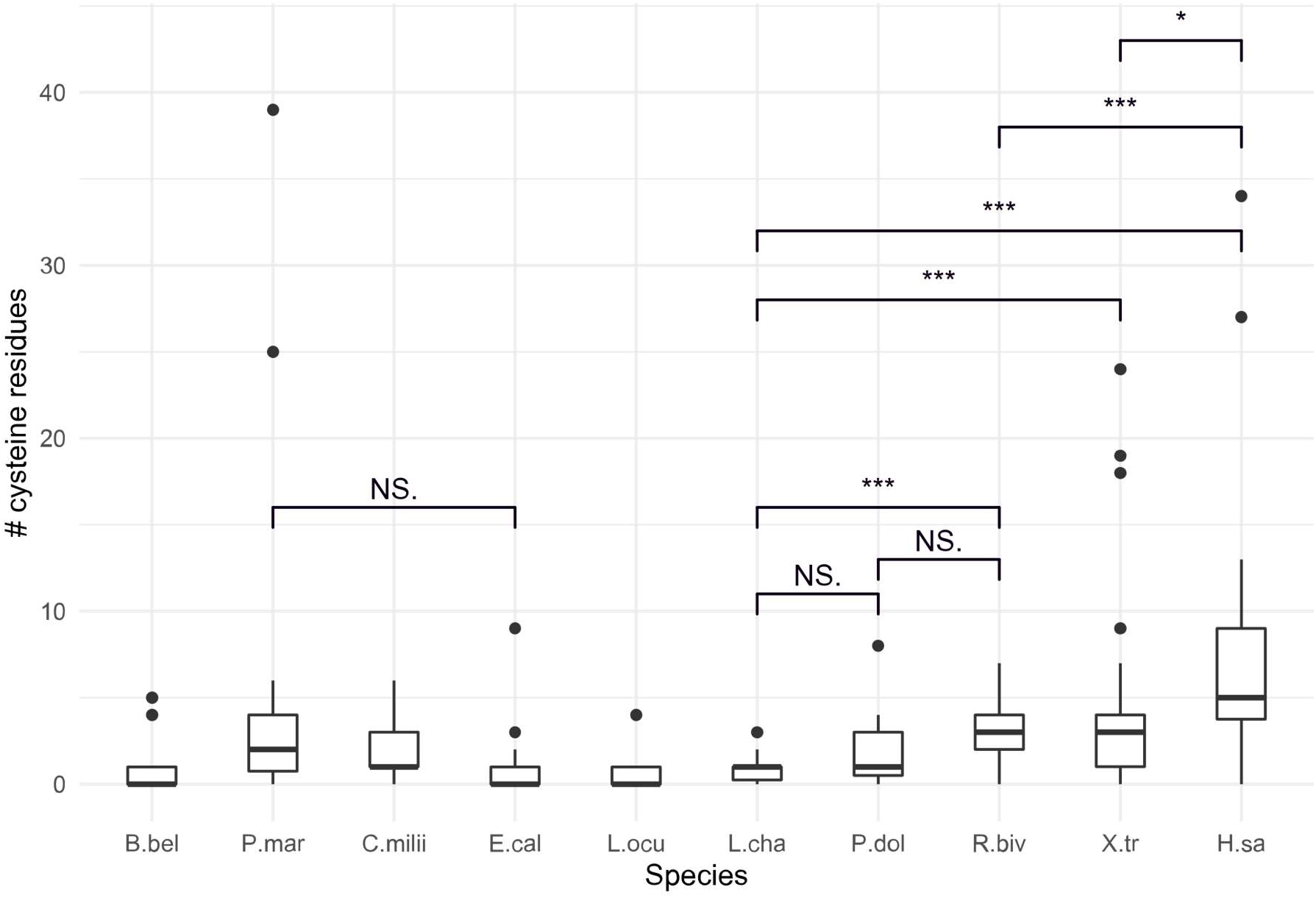
Boxplot of number of cysteine residues in each keratin protein. The species included in the analysis are the following: amphioxus (*Branchiostoma belcheri*; B.bel), lamprey (*Petromyzon marinus*; P.mar), elephant shark (C. *milii*; C.milli), reedfish (E. *calabaricus*; E.cal), spotted gar (L. *oculatus*; L.ocu), coelacanth (L. *chalumnae*; L.cha), lungfish (*P. dolloi*; P.dol), caecilian (R. *bivittatum*; R.bivi), western clawed frog (*X. tropicalis*; X.tr) and human (*Homo sapiens*; H.sa). Note that hair keratins of human were not included in this analysis. t-test was calculated between some species. *, ***: P-values significant at <0.05, <0.001 levels, respectively.

**Supplementary Fig. S3.**
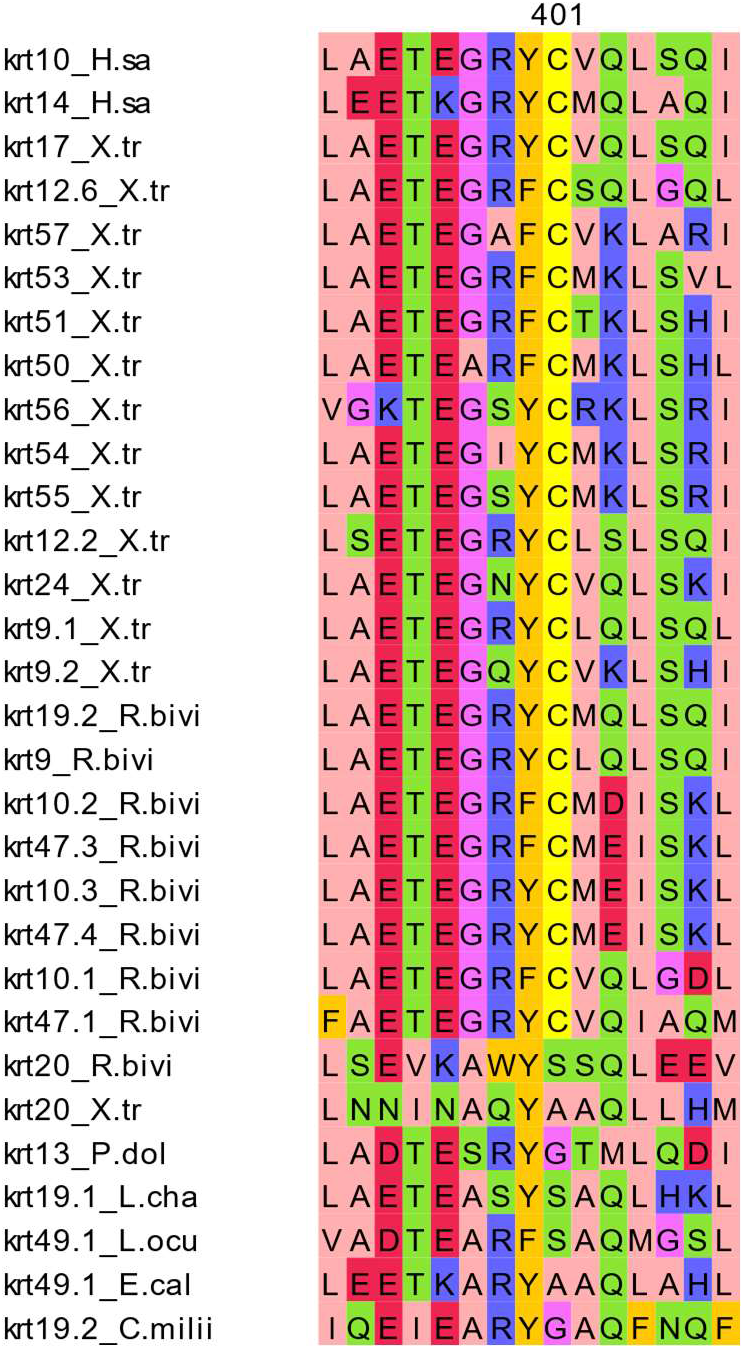
Alignment of partial amino acid sequences of type I keratins with and without the conserved cysteine residue. The species included in the alignment are the following: elephant shark (*C. milii*; C.milli), reedfish (*E. calabaricus*; E.cal), spotted gar (*L. oculatus*; L.ocu), coelacanth (L. *chalumnae*; L.cha), lungfish (*P. dolloi*; P.dol), caecilian (*R. bivittatum*; R.bivi), western clawed frog (*X. tropicalis*; X.tr) and human (*Homo sapiens*; H.sa). Representative keratins without the conserved cysteine residues were also included in the alignment. The number ‘401’ above the alignment indicate the 401^st^ residue of human *krt10*.

**Supplementary File S1.**

FASTA format files for the amino acid sequences of the keratin gene used in the phylogenetic analyses in Fig. 2.

**Supplementary File S2.**

FASTA format files for the nucleotide sequences of the keratin gene used in the selection analyses in Table 1.

**Supplementary File S3.**

FASTA format files for the amino acid sequences of the keratin gene used in the phylogenetic analysis in Supplementary Fig. S1.

**Supplementary Table S1.**

Keratin gene information (i.e., ID, location, and orientation) of vertebrates and amphioxus analyzed in this study.

**Supplementary Table S2.**

Summary of the expression analyses for the RNA-seq data of *Rhinatrema bivittatum* and *Xenopus tropicalis*.

## Notes

### Competing Interest Statement

The authors have declared no competing interest.

### Summary of Updates

We added information about cysteine residues of keratins (Section 4.6; Fig. S2-S3). We changed the name of some keratin genes (e.g. krt14 to krt48) to avoid confusion in referring krt14 in mammals. We added discussion about the significance of keratin clusters in fish.

